# Investigating the repeatability and behavioral relationships of acuity, contrast sensitivity, form, and motion perception measurements using a novel tablet-based vision test tool

**DOI:** 10.1101/2025.06.09.658584

**Authors:** Jonathan Green, Jan Skerswetat, Peter J. Bex, Gunnar Schmidtmann

**Affiliations:** School of Health Professions, University of Plymouth, Plymouth, United Kingdom; Department of Psychology, Northeastern University, 360 Huntington Ave, Boston, Massachusetts, 02115, United States of America; Department of Ophthalmology, University of California, 850 Health Sciences Road, Irvine, California, 9269, United States of America

**Keywords:** visual acuity, contrast sensitivity, form perception, motion perception, psychophysical methods, response-adaptive, self-administered, tablet-based, vision test, angular indication measurement, foraging interactive D-prime, repeatability

## Abstract

Visual function tests are important in basic and clinical vision research but are typically limited to very few aspects of human vision, coarse diagnostic resolution, and require an administrator. Recently, the generalizable, response-adaptive, self-administered Angular Indication Measurement (AIM) and Foraging Interactive D-prime (FInD) methods were developed to assess vision across different visual functions. The AIM and FInD paradigms show a range of visual stimuli per display (4×4 stimuli) spanning ±2σ around an adaptively estimated perceptual threshold across multiple displays. Here, we investigated the repeatability and behavioral relationships of the AIM and FInD paradigms for near visual acuity, contrast sensitivity function (CSF), form, and motion coherence threshold measurements using a novel tablet-based vision test tool. 31 healthy participants were recruited and completed two repetitions of each experiment in random order. Bland-Altman analyses were performed to calculate the Coefficient of Repeatability (precision) and Mean Bias (accuracy). Linear regressions and hierarchical cluster analysis were used to investigate the relationship between outcome parameters. Results show that AIM Form coherence and FInD Form horizontal coherence showed significant retest bias; all other tests were bias-free. Cluster analysis revealed overall clustering of CSF, form and motion outcomes. We further show significant correlations within CSF and between motion coherence outcomes but few significant correlations between form coherence outcomes. AIM and FInD near vision tests are generalizable across multiple visual functions and are precise and reliable. Most functions tested were bias-free. CSF, form, and motion outcomes clustered together, and CSF and motion outcomes correlated with one another. The combination of a generalizable, response-adaptive, and self-administered approach may be a suitable set of tests for basic science and clinical use cases.

## Introduction

Identification and measurement of visual dysfunction is a global challenge. In excess of 1 billion people have visual impairments that could have been or are yet to be addressed (World Health Organisation (WHO), 2019; Bourne *et al*., 2019) with the estimated associated loss of productivity valued at US$ 410.7 billion (WHO, 2023). This challenge is exacerbated by the disproportionate prevalence of visual deficiency in lower and middle-income countries (90%) (WHO, 2023; Rizzo *et al*., 2020; WHO, 2022) and in higher income countries, amongst those from ethnic minorities, with lower socio-economic status, lower educational attainment (Cumberland *et al*. 2016; Elam *et al*. 2022), and poorer general and mental health (Cumberland *et a*l. 2016).

Ageing is also strongly associated with increased risk for many visual conditions (Killeen *et al*., 2023; Martinez-Roda *et al*., 2016) - including subtle and under-recognised presentations, such as those often associated with mild traumatic brain injury (Rasdall *et al*., 2024) (over 1-million annually in the US alone). Globally, the number of people aged over 65 is predicted to double over the next quarter of a century (United Nations, 2022), representing a substantial generator of pressure on often already overstretched health services.

Innovative digital technologies offer a range of opportunities to mitigate such challenges (Chen, Ding & Wang, 2023). Many digital vision assessment tools are available, some of which can now be operated using conventional portable devices such as laptop and tablet computers, or even smartphones (Bano, Wolffsohn & Sheppard, 2024). Advances in functionality support rapid, non-specialist, point-of care and domiciliary assessment, as well as self-administration and remote operation (telemedicine, Samanta *et al*., 2023). These technologies promise to drive more efficient assessment and monitoring, at earlier disease stages, reducing inconvenience and expense, and increasing access to underserved patient groups (Khurana *et al*., 2021). As yet, these novel systems generally target a limited range of visual functions – and many are yet to be subjected to robust, peer-reviewed validation of their accuracy and reliability (Bano, Wolffsohn & Sheppard, 2024).

This study focuses on new ‘Perzeption’ suites of computer-based visual assessment tools which apply novel psychophysical paradigms and algorithms; namely the Angular Indication Measurement (AIM) (Skerswetat *et al*., 2024a; Neupane, Skerswetat & Bex, 2024) and the Foraging Interactive D-prime (FInD) (Bex & Skerswetat, 2021; He, Bex & Skerswetat, 2023; Skerswetat *et al*., 2024b). Tests employing these generalisable, self-administered, multi-dimensional (spatial/temporal frequency/contrast), and response-adaptive (personalised) psychophysical methods have been developed to measure more than 15 visual functions (Skerswetat *et al*., 2024b) including visual acuity with and without glare source (Skerswetat, Boruta & Bex, 2022), colour (He, Bex & Skerswetat, 2023) and depth perception (Neupane, Skerswetat & Bex, 2024), spatio- and spatio-temporal contrast sensitivity (Skerswetat *et al*., 2024b), threshold versus contrast suprathreshold perception (Bex & Skerswetat, 2021; Bex & Skerswetat, 2023), form and motion perception (Bex & Skerswetat, 2021; Skerswetat *et al*., 2024b; Merabet *et al*., 2023), and face discrimination (Walter & Bex, 2024). Advantages of these methods compared to traditional clinical tests and current laboratory computer-based methods include flexible testing ranges, fine outcome resolution and improved detection of subtle changes, a bias-free test protocol due to randomised stimulus orientations and variables that eliminate memorisation risks, rapid testing through adaptive algorithms and instant result generation, reduction in administrative burden and time due to self-administration, and a generalizable method to measure multiple visual functions using the same principal, i.e., orientation judgement for AIM and signal presence detection for FInD.

The AIM method can be applied with different stimulus and response formats, generating orientation judgements. Psychometric functions are fit to continuous orientation report errors (as a function of stimulus intensity) (Skerswetat *et al*., 2024a). The FInD method measures the identification of cells containing signal stimuli of varying intensities, amongst null stimuli cells. Thresholds are estimated from *d’* analysis of hits, misses, false alarms, and correct rejection responses (Skerswetat *et al*., 2024b).

Previous studies have supported the refinement and administration of many of these assessment tools. One study demonstrated the systematic application of a battery of eleven tests, across both paradigms, for the broad visual assessment of 21 research participants with visual dysfunction (people with albinism), compared with 21 typically sighted controls (Skerswetat *et al*., 2024b). Anticipated variance between groups was observed – and the incorporation of testing pertinent to different levels of visual processing provided insight into a lack of differentiation for some mid-level functioning (pattern recognition, and radial and circular motion coherence). The FInD global motion coherence test has also been demonstrated to identify an anticipated significant reduction in performance amongst a cohort of 20 young people with cerebral visual impairment, compared with 30 controls (mean difference 32.34%) - concurrently providing evidence of usability by neurodiverse participants and for the targeted assessment of cerebral visual processing (Merabet *et al*., 2023). Participants were able to use both form and motion tests, taking an average of around 6 minutes to complete both.

In a study of 22 young adults with induced defocus and astigmatic blur, results from the AIM visual acuity (VA) test demonstrated comparable sensitivity to conventional Early Treatment Diabetic Retinopathy Study (ETDRS) letter chart assessment and comparable repeatability when the AIM test includes three adaptive steps (mean VA test retest bias 0.03 and 0.02, respectively), whilst providing additional personalised information regarding minimal error level (noise), bias and slope from a psychometric function fit (Skerswetat *et al*., 2024a). Mean AIM VA test duration was only around 30 seconds. Elsewhere, a study of 24 children (including 11 amblyopes) demonstrated the usability of AIM VA for children as young as 5 years, taking an average of around 1.5 minutes (Ahmed *et al*., 2023). The results demonstrated good agreement with Snellen and Rosenbaum chart use (ANOVA group effect = 0.612) and identified interocular VA differences.

The AIM Contrast Sensitivity Function (CSF) test was appraised in a study of 10 participants (8 typically sighted, 1 amblyopic and 1 Ortho-K contact lens wearer with defocus blur) (Skerswetat & Bex, 2022). Between-eye results were similar, and less than for both eyes together, consistent with binocular contrast summation.

Anticipated interocular difference and overall performance reduction was observed for the respective atypically sighted participants. CSF testing was generally completed within 2 minutes. Elsewhere, results of FInD CSF and Threshold versus Contrast functions (TvC) [stereoacuity, spatial coherence and motion coherence] tests were found to be comparable to the results of equivalent standard temporal two-alternative forced choice (2-AFC) paradigms for up to 14 participants (Bex & Skerswetat, 2021). The FInD tests were significantly quicker to administer, and no significant test retest difference or *bias* were detected.

Whilst this is a test retest study, it is of note that there is also an accumulation of evidence regarding the performance of other tests in the Perzeption batteries compared to established measures. The AIM Threshold versus Contrast (TvC) and Equivalent Noise (EN) assessments for form, motion and color have been demonstrated to be comparable to two-alternative forced choice (2-AFC) tests (Bex & Skerswetat, 2023). Elsewhere, both FInD and AIM tests for stereopsis reliably detected deficits compared to 2-AFC and Titmus and Randot clinical tests, although stereo thresholds were 1.3-1.6 times higher for FInD and 2-AFC than for the clinical tests (Neupane, Skerswetat & Bex, 2024). Regarding color vision assessment, FInd and AIM tests agreed well with Cambridge Color Test (CCT) thresholds and Mollon-Reffin scores for the detection of color vision deficiency (CVD). AIM and FInD performance slightly reduced for CVD categorisation, but their combined results were comparable to the other tests (Arthur *et al*., 2025). The FInD test reliably categorised CVD compared to Hardy-Rand-Rittler and Farnsworth-Munsell 100 hue tests, accepting that even the results of these two other methods were not entirely uniform (He, Bex & Skerswetat, 2023) and in another study the AIM test was demonstrated to have good agreement with anomaloscope readings for CVD detection (He, Skerswetat & Bex, 2023).

Performance across visual perception domains does not appear to be highly correlated in healthy adults, suggesting multifactorial determinants of normal visual function (Tulver, 2019). Attempts to map consistent relationships between potentially associated sub-groups of functions have also, as yet, only yielded modest correlations (Bosten *et al*., 2017a; Cappe et al., 2014; Herzog *et al*., 2025; Shakiri *et al*., 2019). Intra-observer variance appears to be stable over time, i.e. consistently deteriorating (Garobbio, Kunchulia & Herzog, 2014), and certain traits may be associated with specific disorders (Bosten *et al*., 2017b). The inherent efficiency and the relative homogeneity of interface, stimulus type, format, etc. of this test battery may prove useful in the advancement of research in this field.

The aim of the current study is to investigate the repeatability and behavioral relationships of the AIM and FInD paradigms for acuity, contrast sensitivity, form, and motion perception measurements implemented as a novel tablet-based vision test tool.

## Methods

### Participants

31 observers (mean age: 39.7, **𝜎**: ±11.7; 13 females) participated in the initial test-retest study. Seven additional observers (mean age: 39, **𝜎**: ±15; 4 females) were recruited for a control experiment (AIM CSF with five displays, see details below). 36 observers were naive as to the purpose of the experiment. All participants had normal or corrected-to-normal visual acuity. To assess whether performance in the visual tests was influenced by Asperger syndrome (i.e., high-functioning autism) or attention deficit hyperactivity disorder (ADHD), all participants completed the Adult ADHD Self-Report Scale (ASRS; Kessler et al., 2005) and the Autism-Spectrum Quotient (AQ; Baron-Cohen et al., 2001) questionnaires (see Table 1 caption for details). Participants also provided information regarding their medical history. One participant reported hemiplegic migraines, three reported past (non-recent) concussions, one reported anxiety, emotional dysregulation, and depression, and another had a diagnosis of dyslexia. None of these conditions were considered sufficient grounds for exclusion. Reported ocular conditions (e.g., myopia, astigmatism) did not affect participant eligibility, as all tests were conducted with appropriate visual correction. Likewise, reported colour vision deficiencies were not considered exclusionary, since none of the tasks involved coloured stimuli.

**Table 1.** Self-reported participant demographic and medical details. Ethnicity: White (UK, Irish) includes English, Welsh, Scottish, Irish, British. Sex: f = female; m = male. ADHD & Autism Thresholds: Adler’s Adult ADHD Self-Report Scale (ASRS-v1.1). Part A (screener) ≥ 4 grey boxes = highly consistent with ADHD; Baron-Cohen’s Autism Spectrum Quotient (AQ): ≥32 = Clinically significant (80% of those diagnosed with autism, but <2% of controls), ≥ 26 = Potential traits

| ID | Age | Ethnicity | Sex | Medical history | Ocular conditions | ASRS | AQ |
| --- | --- | --- | --- | --- | --- | --- | --- |
| 1 | 48 | White (UK, Irish) | m |  | Myopia | 0 | 20 |
| 2 | 44 | White (UK, Irish) | m |  |  | 0 | 14 |
| 3 | 48 | White (UK, Irish) | m | Hemiplegic migraine | Astigmatism | 1 | 23 |
| 4 | 22 | Black | m |  |  | 0 | 11 |
| 5 | 44 | White (UK, Irish) | f |  |  | 0 | 7 |
| 6 | 35 | White (UK, Irish) | f |  |  | 2 | 15 |
| 7 | 42 | White (UK, Irish) | m | Concussion |  | 1 | 18 |
| 8 | 49 | White (UK, Irish) | m |  |  | 0 | 23 |
| 9 | 39 | White (UK, Irish) | m |  |  | 0 | 11 |
| 10 | 29 | White (UK, Irish) | f | Anxiety, Emotional dysregulation, Depression |  | 3 | 10 |
| 11 | 28 | White (UK, Irish) | m |  |  | 2 | 22 |
| 12 | 23 | Asian | m |  | Color-Vision-Defect | 0 | 12 |
| 13 | 20 | Black | f |  |  | 0 | 17 |
| 14 | 28 | White (UK, Irish) | f |  |  | 2 | 10 |
| 15 | 28 | White (UK, Irish) | m |  |  | 5 | 39 |
| 16 | 48 | White (UK, Irish) | m |  |  | 2 | 21 |
| 17 | 26 | Asian | m |  |  | 2 | 25 |
| 18 | 32 | White (UK, Irish) | m | Concussion |  | 3 | 28 |
| 19 | 29 | White (UK, Irish) | m | Concussion |  | 2 | 21 |
| 20 | 19 | White (UK, Irish) | f |  |  | 2 | 12 |
| 21 | 57 | White (UK, Irish) | f |  |  | 0 | 11 |
| 22 | 54 | White (any other) | f |  |  | 0 | 10 |
| 23 | 48 | White (UK, Irish) | f | Dyslexia |  | 2 | 14 |
| 24 | 43 | White (UK, Irish) | m |  |  | 2 | 20 |
| 25 | 58 | White (UK, Irish) | f |  |  | 1 | 15 |
| 26 | 35 | White (UK, Irish) | m |  |  | 0 | 8 |
| 27 | 50 | White (UK, Irish) | f |  | Astigmatism | 0 | 15 |
| 28 | 55 | White (UK, Irish) | f |  |  | 2 | 24 |
| 29 | 46 | White (other) | m |  |  | 0 | 13 |
| 30 | 51 | White (UK, Irish) | f |  |  | 0 | 10 |
| 31 | 40 | mixed | m |  |  | 0 | 30 |

Informed, written consent was obtained from each observer, and the study was approved by the University of Plymouth ethics committee. All experiments were conducted in accordance with the Declaration of Helsinki.

### Stimuli and psychophysical procedure

Each display chart for each test contained a 4×4 set of cells. All tests used three charts per experimental run, including the first initial chart and two adaptive steps for all tests. The range of stimulus test levels on each display spanned ±2σ around an adaptively estimated perceptual threshold across displays (e.g., Skerswetat *et al*., 2024b; Skerswetat, Boruta & Bex, 2022). For form and motion perception testing, the default display mode, which presents all stimuli simultaneously and also the hidden cell approach, which shows only the stimulus over which the user positions the cursor (Merabet *et al*., 2023) was used. Orientations or directions of stimuli within cells were randomized for all AIM tests. Stimulus examples are shown in Figure 1.

**Figure 1.**
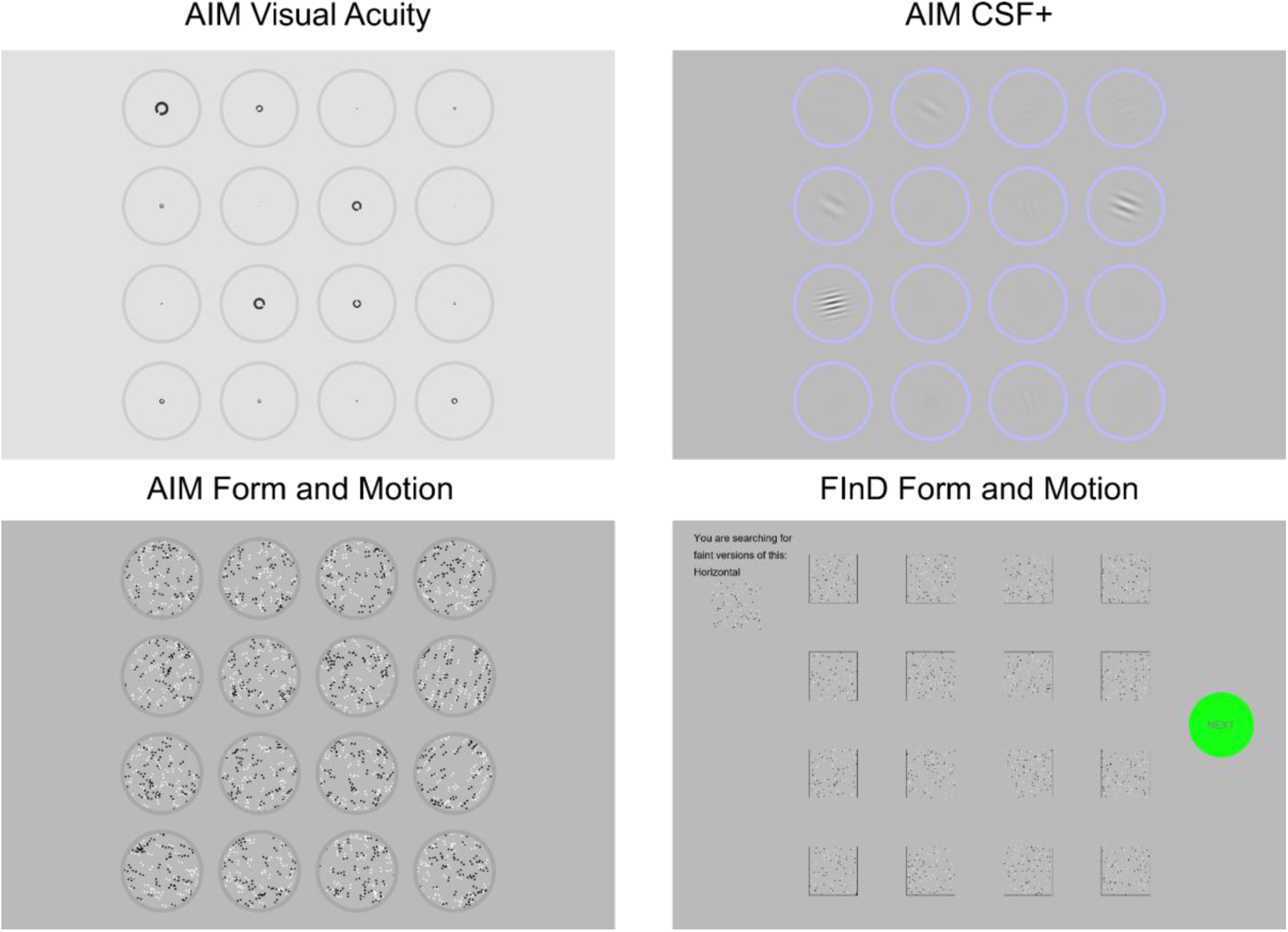
The images show charts containing stimulus examples for the AIM Visual Acuity (VA), AIM CSF+, AIM Form and Motion Coherence and FInD Form and Motion Coherence tests as seen by the observers on the tablet.

### Angular Indication Measurement (AIM)

Participants indicated the perceived orientation of all 16 stimuli on one chart by clicking on each surround ring (see Figures 1 and 2) or making their best guess if they were not sure. The testing time was unlimited, and correction of indications were permitted prior proceeding to the next chart by clicking on a ‘Next’ button that became available only once responses had been made to all 16 stimuli (see bottom right in Figure 1). The response error was calculated as the absolute angular difference between target orientation and indicated orientation. After the final chart was completed, a cumulative Gaussian function was fit to the absolute angular error data as a function of stimulus intensity, namely optotype size (AIM-VA), grating contrast (AIM CSF+), translational form or motion coherence (AIM-Form and AIM-Motion) from all charts, defined as

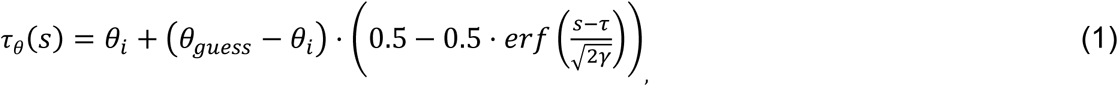

where *θ_guess_* denotes the orientation error for a guess response (the absolute orientation error of random responses for stimuli is uniformly distributed with a mean of 90° for Landolt C optotypes, translation form and motion coherence). For Gabors during AIM-CSF2d+, the rotational symmetry *θ_guess_* is 45°. *s* denotes the signal intensity, *𝜏* a sensitivity threshold, *θ_i_* is internal angular uncertainty, the absolute angular report error for the most visible stimulus, which we refer to as a *noise* parameter as it is due to some of the ex- and internal noise sources, and *γ* is the slope of the psychometric function.

**Figure 2:**
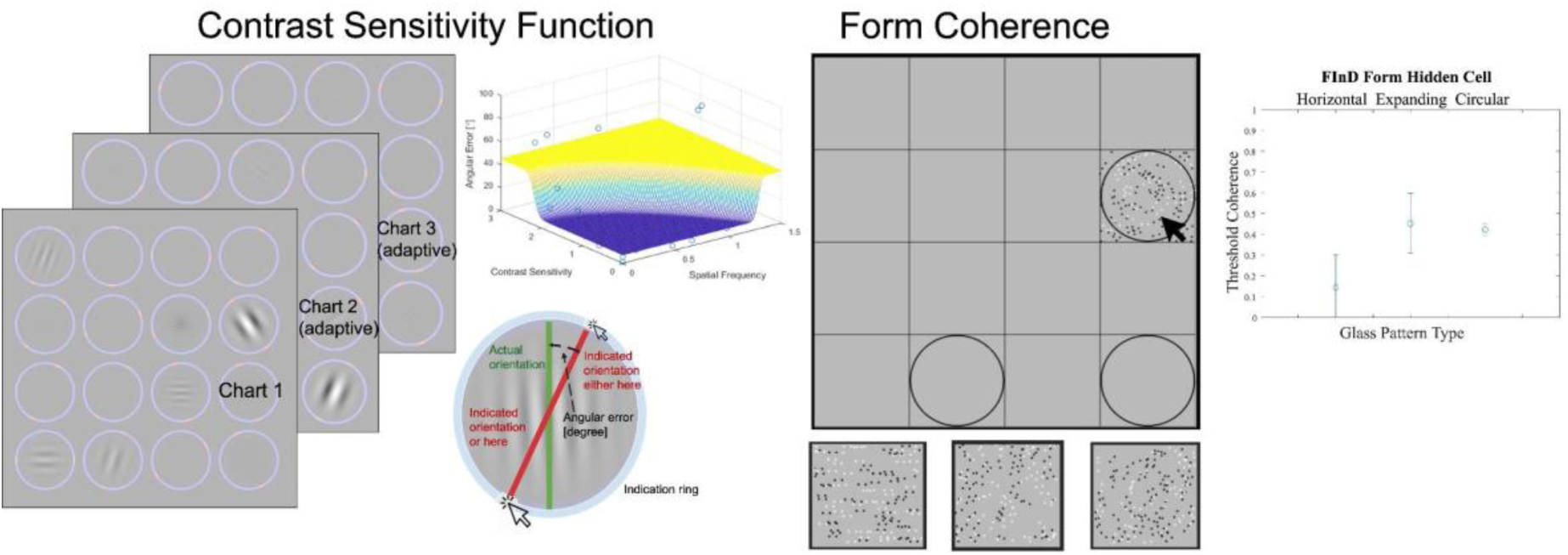
Schematic illustration of the AIM (left) and FInD (right) methods. Left: The AIM method is exemplified by the AIM-CSF 2D+ paradigm. The initial chart showed Gabor patches with a range of different spatial frequencies and contrasts. Participants then indicate the perceived orientation of each Gabor or make their best guess before continuing to the next chart by pressing a green “Next button” that appears after all 16 indications are made (not shown here). Subsequent charts adaptively homing in around the CSF. After completing the final chart, the *log* parabola function was fit to the absolute report errors to estimate the area under the log CSF and CSF-acuity. The difference between the actual Gabor orientation (green line) and indication orientation (red line) are shown underneath the example fit. The right illustration shows the FInD Form using the hidden cell method. The initial chart contains 16 cells, 9-12 of them contain a signal (e.g., circular Glass patterns) of varying intensities (randomly positioned within the chart), and all other cells contain noise. The participant uses a computer mouse to hover over each cell to make it appear, judges whether it is a signal and makes their indication before moving to the next cell. The participant was instructed to “zigzag” over each cell before clicking on the “Next” button, which contained a different form-coherence (horizontal, expanding, circular in random order). Once the first initial chart for each form coherence type was completed, the next charts adaptively converged on ±2 standard deviation around each form pattern threshold and then repeated until three charts per form coherence type were completed. The psychometric functions and their 95% confidence intervals (CI) were fit to each form coherence type. An example outcome for FInD Form thresholds and 95% CIs is depicted on the right.

#### AIM Acuity

Landolt C optotypes were presented within each indication ring (diameter, Ø: 5°) with an inter-cell distance of 1°. Stimulus contrast was 0.97. The minimum range of stimuli sizes was set to 0.20 logMAR. The minimum visual acuity was kept at the minimum renderable size for −0.05 logMAR and maximum logMAR starting value was set to 1.00 logMAR. More details are published elsewhere (Skerswetat *et al*., 2024a).

#### AIM Form Coherence (with and without Hidden Cells)

Glass Pattern (Glass, 1969; Glass & Perez, 1973) coherence for translational form coherence was measured with 100 dipoles (50 black, 50 white; cell Ø: 5°; dot Ø=0.1°; intra-dipole spacing=0.2°).

#### AIM Motion Coherence (with and without Hidden Cells)

Motion coherence for translational global dot motion was measured. Each cell (Ø: 5°) contained 100 black or white dots (Ø=0.1°, lifetime=100ms with random starting age, speed=2°/sec, dot contrast: 1.0). Stimulus examples are presented in the supplementary AIM_Motion.mp4 video file.

#### AIM CSF 2D+

Gabors (σ: 0.5°) were presented within each indication ring (Ø: 5°) with an inter-cell distance of 1°.

One initial and two adaptive charts were presented and the task was to click on the ring surrounding each Gabor to indicate the orientation of the grating or to make a guess (Figure 2). We assessed the CSF acuity (high spatial frequency cutoff) and the area under the *log*CSF (AULCSF). In order to increase the greyscale resolution of the 8-bit monitor, we applied the noisy-bit binomial sampling method (Allard & Faubert,2008).

The angular report errors were fit with an asymmetric *log* parabola to estimate the CSF.

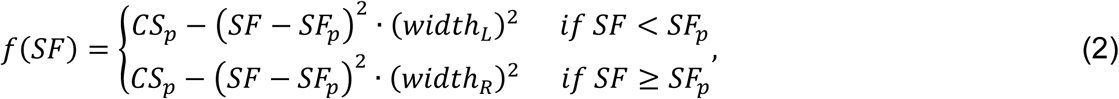

The variable *CSp* denotes the peak contrast sensitivity, *SFp* the peak spatial frequency, and *width_L_* and *width_R_* are the lower and upper bandwidths (Watson & Ahumada, 2005).

#### AIM specific feedback

Auditory feedback was provided after each chart based on a statistical probability analysis of per-chart report errors. If the mean error report was indistinguishable from random responses (i.e., 90° error for C optotypes and motion directions, 45° for gratings and Glass patterns), the application altered the user to report the orientation carefully.

### Foraging Interactive D-prime (FInD)

The right panel in Figure 2 illustrates the FInD method. The task for the participants was to select cells for each chart that contained a stimulus. No time-constraint was given. Selected cells could be unselected within each chart and were only locked after pressing the green “Next” button to initiate the next display (see bottom right in Figure 1). The response in each cell was categorised as either Hit, Miss, False Alarm, or Correct Rejection, to calculate *d’,* and the probability of a Yes response as a function of signal intensity was calculated according to

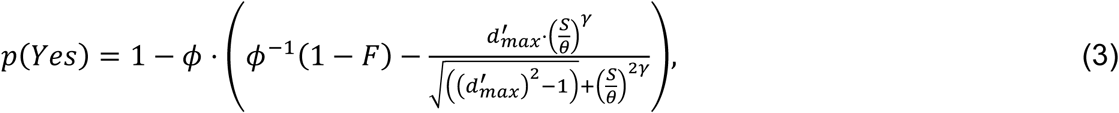

where *p(Yes)* is the probability of a Yes response, *ϕ* is the normal cumulative distribution function, and *F* is the false alarm rate, *S* denotes the stimulus intensity, *θ* the threshold, *d’_max_* is the saturating value of *d’* (fixed at 5), and *γ* is the slope. The fit to data from all completed charts was used to select the individualized range (i.e., *d’* = 0.1 to 4.5) of stimulus intensities (namely static Glass (Glass & Perez,1973) or motion coherence patterns) for subsequent charts.

#### FInD Form (with and without Hidden Cells)

Glass Pattern coherence (Merabet *et al*., 2023; Glass, 1969; Glass & Perez, 1973) for translational, circular, and expanding (radial) form coherence was measured with 100 dipoles (50 black, 50 white; cell Ø: 3×3° squares; dot Ø=0.05°; inter-dot spacing=0.2°). The inter-cell distance was 3°.

#### FInD Motion (with and without Hidden Cells)

Global motion coherence (Merabet *et al*., 2023) for translational, circular, and expanding form coherence was measured with 100 dipoles (50 black, 50 white; dot contrast: 1.0; cell Ø: 3*3° squares; dot Ø=0.05°; speed=1°/sec, lifetime=100ms with random starting age). The inter-cell distance was 3°.

### Correlation and cluster analysis

Python, including the Seaborn toolbox (Waskom, 2021) was used to perform and visualize the linear correlation and hierarchical cluster analysis. Data were normalized using z-score normalization. The marginal cluster trees show a hierarchical cluster analysis using Ward’s Minimum Variance Clustering Method based on the Euclidean distance between objects (i.e., test results). Linear correlations using Kendall coefficients were used.

### Apparatus

Stimuli were presented on a Microsoft Surface Pro 8 (Core^TM^ i5 CPU 8GB, 256GB Go RAM) with display dimensions of 27.4 x 18.3 cm. The display’s spatial resolution was 2880 x 1920 pixels with a 60Hz frame rate. All experiments were carried out at a viewing distance of 40 cm. The experimenter monitored the viewing distance throughout the task and provided feedback if the viewing position was changed. Note that while participant responses are therefore self-administered and data analysis was automated, monitoring of viewing distance therefore involved supervision by the examiner. At this distance, one pixel subtends 0.0136° visual angle. The screen was gamma-corrected using data color Spyder (Datacolor, Lucerne, Switzerland) photometer. The input for the AIM tests was via the tablet’s touchscreen. The AIM hidden cell and FinD paradigm reports were made using a common computer mouse.

### Procedure

Repeatability of near visual acuity, contrast sensitivity function (CSF), and form & motion coherence tasks were deployed on a tablet (see above). Observations were made under binocular viewing conditions. The test order was randomised, each experiment included a test and a retest session and took in total ∼90 min.

## Results

### Psychophysical outcomes

The PerZeption application generated automatically outcomes as described below.

**AIM:** A cumulative Gaussian function (equation 1) was fit to the data of the AIM Visual Acuity, AIM Form Coherence, and AIM Motion Coherence tests (Neupane *et al*., 2024; Skerswetat *et al*., 2024b). Threshold, slope, and minimum report error parameter were calculated. For the AIM CSF 2D+, the area under the log CSF (AULCSF) and the high spatial frequency cut off, or CSF Acuity were analysed.

**FInD:** Response of each cell was classified as either Hit, Miss, False Alarm, or Correct Rejection, to calculate *d’* as a function of motion or form coherence. The probability of a Yes response as a function of signal intensity was subsequently calculated (equation 3, Merabet *et al*., 2023). Threshold (*d’*=1), slope, and false alarm rate parameters were calculated.

### Bland-Altman Analysis

Test-retest reliability for all tests was assessed using Bland-Altman (Altman & Bland, 1983) plots using MATLAB (R2024b, Matworks). Figure A1 in the appendix shows an example (see Supplementary Material for all Bland-Altman plots).

We subsequently calculated the Coefficient of Repeatability (precision, *CoR*) according to

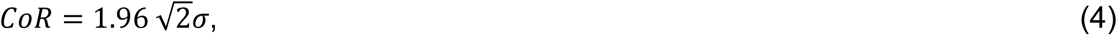

where **𝜎** refers to the within-subject standard deviation. The mean bias (<u>𝑥</u>_𝑏𝑖𝑎𝑠_ , accuracy) in Table 2 was calculated as

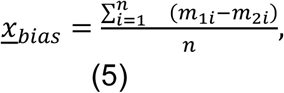

where *m_1_* and *m_2_* denote the test and retest measurements, respectively. One-sample t-tests were conducted to assess whether the bias is statistically significant. Test-retest data (*m_1_*, *m_2_*), *CoR*, mean *Bias* and *p*-values are summarized in Table 2, where 𝑝 ≤ 0.05 is considered statistically significant and indicated by asterisks.

**Table 2.**
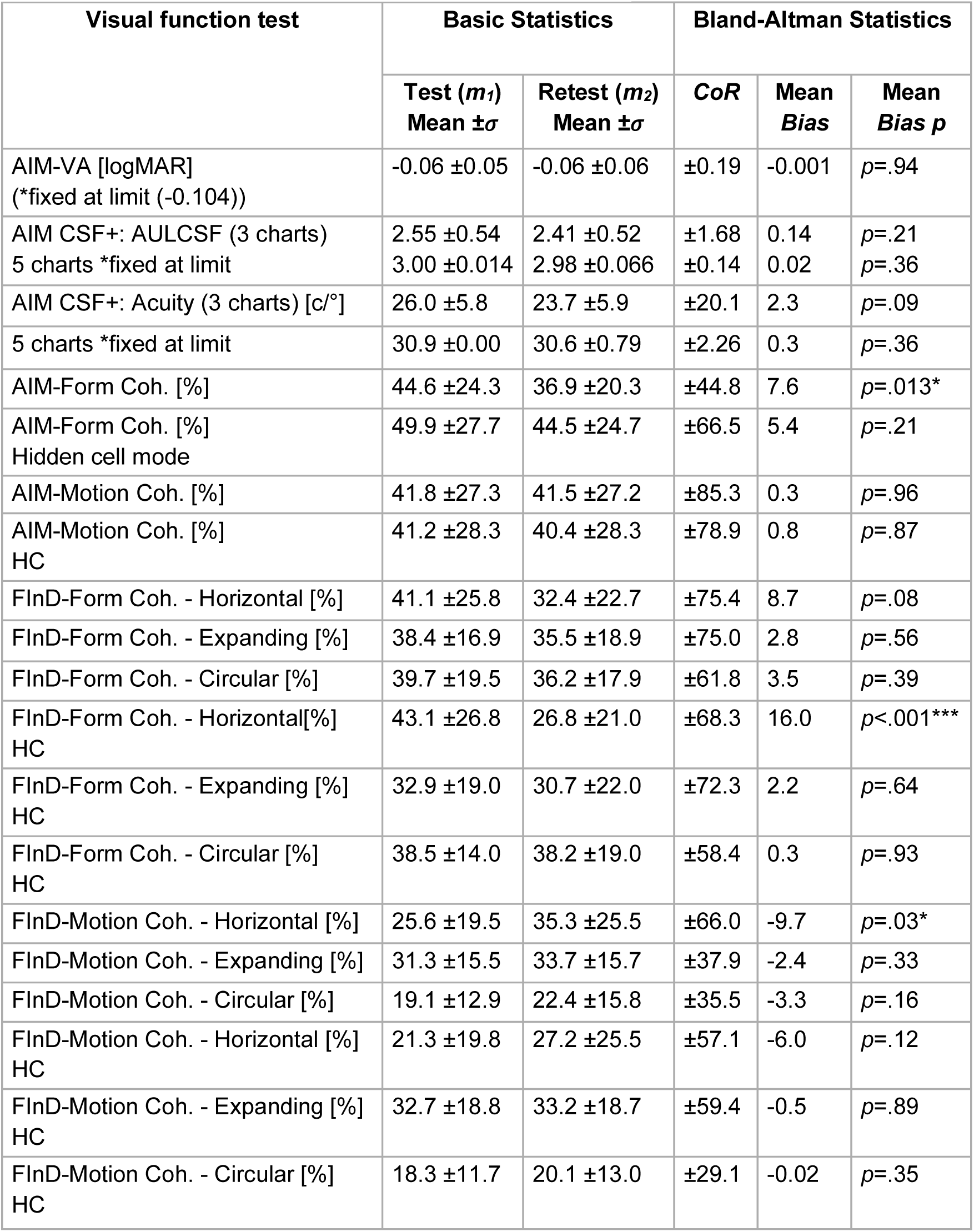
Test Retest results for threshold values (Coh. = coherence threshold, HC = hidden cell mode, 𝜎 = standard deviation, *CoR* = coefficient of repeatability).

| Visual function test | Basic Statistics |  | Bland-Altman Statistics |  |  |
| --- | --- | --- | --- | --- | --- |
| | Test ($m_1$)<br>Mean $\pm \sigma$ | Retest ($m_2$)<br>Mean $\pm \sigma$ | CoR | Mean Bias | Mean Bias $p$ |
| AIM-VA [logMAR]<br>(*fixed at limit (-0.104)) | -0.06 ± 0.05 | -0.06 ± 0.06 | ±0.19 | -0.001 | $p=.94$ |
| AIM CSF+: AULCSF (3 charts)<br>5 charts *fixed at limit | 2.55 ± 0.54<br>3.00 ± 0.014 | 2.41 ± 0.52<br>2.98 ± 0.066 | ±1.68<br>±0.14 | 0.14<br>0.02 | $p=.21$<br>$p=.36$ |
| AIM CSF+: Acuity (3 charts) [c/°]<br>5 charts *fixed at limit | 26.0 ± 5.8<br>30.9 ± 0.00 | 23.7 ± 5.9<br>30.6 ± 0.79 | ±20.1<br>±2.26 | 2.3<br>0.3 | $p=.09$<br>$p=.36$ |
| AIM-Form Coh. [%] | 44.6 ± 24.3 | 36.9 ± 20.3 | ±44.8 | 7.6 | $p=.013^*$ |
| AIM-Form Coh. [%]<br>Hidden cell mode | 49.9 ± 27.7 | 44.5 ± 24.7 | ±66.5 | 5.4 | $p=.21$ |
| AIM-Motion Coh. [%] | 41.8 ± 27.3 | 41.5 ± 27.2 | ±85.3 | 0.3 | $p=.96$ |
| AIM-Motion Coh. [%]<br>HC | 41.2 ± 28.3 | 40.4 ± 28.3 | ±78.9 | 0.8 | $p=.87$ |
| FInD-Form Coh. - Horizontal [%] | 41.1 ± 25.8 | 32.4 ± 22.7 | ±75.4 | 8.7 | $p=.08$ |
| FInD-Form Coh. - Expanding [%] | 38.4 ± 16.9 | 35.5 ± 18.9 | ±75.0 | 2.8 | $p=.56$ |
| FInD-Form Coh. - Circular [%] | 39.7 ± 19.5 | 36.2 ± 17.9 | ±61.8 | 3.5 | $p=.39$ |
| FInD-Form Coh. - Horizontal [%]<br>HC | 43.1 ± 26.8 | 26.8 ± 21.0 | ±68.3 | 16.0 | $p<.001^{***}$ |
| FInD-Form Coh. - Expanding [%]<br>HC | 32.9 ± 19.0 | 30.7 ± 22.0 | ±72.3 | 2.2 | $p=.64$ |
| FInD-Form Coh. - Circular [%]<br>HC | 38.5 ± 14.0 | 38.2 ± 19.0 | ±58.4 | 0.3 | $p=.93$ |
| FInD-Motion Coh. - Horizontal [%] | 25.6 ± 19.5 | 35.3 ± 25.5 | ±66.0 | -9.7 | $p=.03^*$ |
| FInD-Motion Coh. - Expanding [%] | 31.3 ± 15.5 | 33.7 ± 15.7 | ±37.9 | -2.4 | $p=.33$ |
| FInD-Motion Coh. - Circular [%] | 19.1 ± 12.9 | 22.4 ± 15.8 | ±35.5 | -3.3 | $p=.16$ |
| FInD-Motion Coh. - Horizontal [%]<br>HC | 21.3 ± 19.8 | 27.2 ± 25.5 | ±57.1 | -6.0 | $p=.12$ |
| FInD-Motion Coh. - Expanding [%]<br>HC | 32.7 ± 18.8 | 33.2 ± 18.7 | ±59.4 | -0.5 | $p=.89$ |
| FInD-Motion Coh. - Circular [%]<br>HC | 18.3 ± 11.7 | 20.1 ± 13.0 | ±29.1 | -0.02 | $p=.35$ |

Results show that AIM Form coherence results and FInD Form horizontal coherence results showed significant retest biases, but all other tests were bias-free.

### Exploratory Correlation and Cluster Analysis

After assessing the test’s repeatability and gaining insights into the normative range of performance within each test, we explored how these performance results relate to one another. Hence, we conducted a linear regression and hierarchical cluster analysis to investigate the relationship between outcome parameters. Kendall’s correlation coefficients between all tests were calculated. The matrix in Figure 3 shows the correlation coefficients between all tests as color-code and value in each cell of the matrix. The warmer colors indicate higher and the colder colours lower correlations (see colour bar). The asterisks indicate whether the correlation was significantly different from zero. This analysis reveals significant correlations within CSF and between motion coherence outcomes but few significant correlations between form coherence outcomes. The marginal cluster trees in Figure 3 show results from a hierarchical cluster analysis using Ward’s Minimum Variance Clustering Method based on the Euclidean distance between objects (i.e., tests).

**Figure 3.**
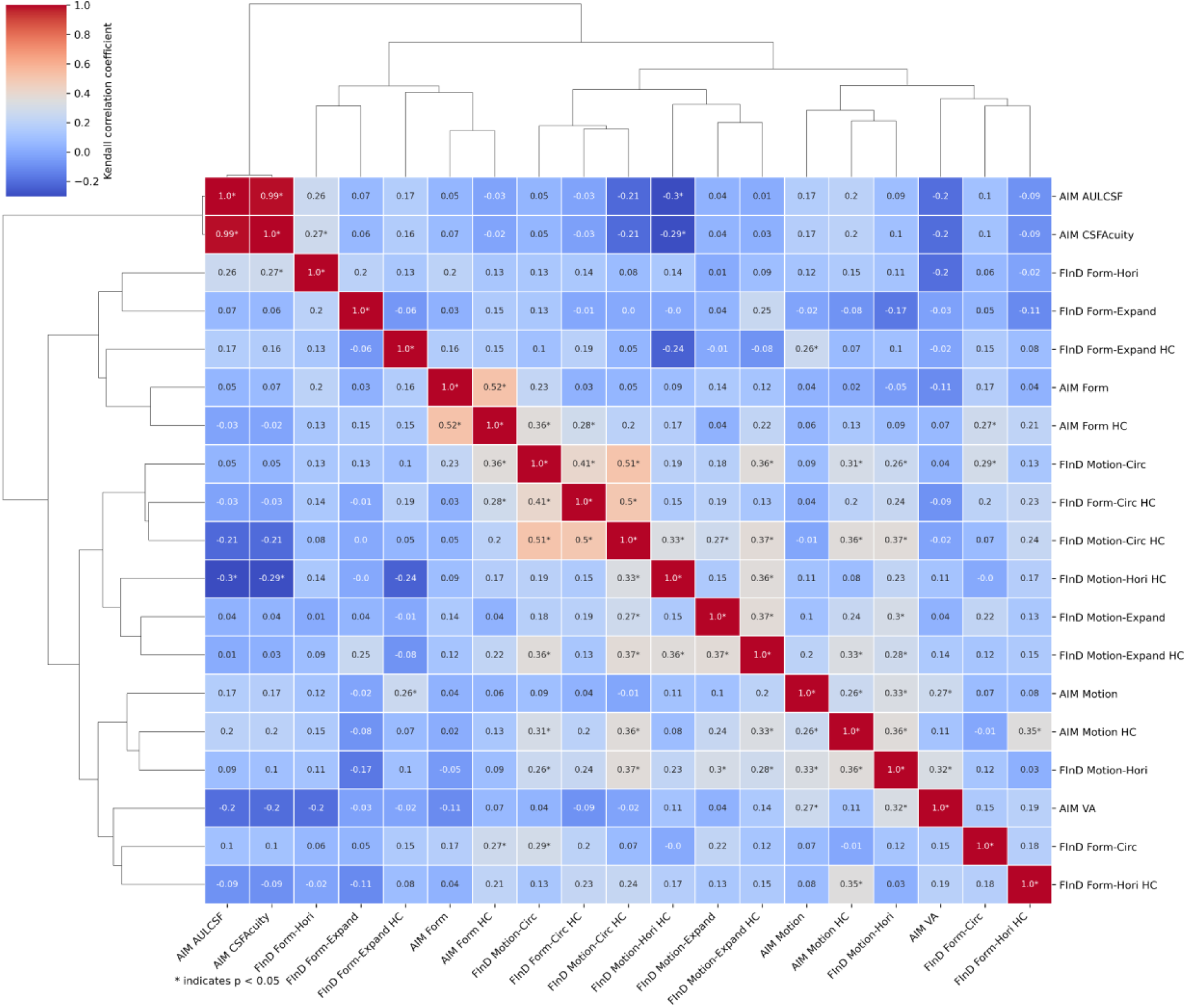
(i) The figure shows Kendall’s correlation coefficients between all tests as color-code and value in each cell of the matrix, where warmer colors indicate higher correlations (see color bar). The asterisks indicate whether the correlation was significantly different from zero. (ii) The marginal cluster trees show results from a hierarchical cluster analysis). Note that the data set includes a dependent set of CSF Acuity and AULCSF data that is derived from the same logparabola fit function in each observer.

This analysis revealed overall clustering of CSF, Form and Motion tests.

### Correlation between performance and ASRS and AQ scores

In future work, we aim to investigate whether performance is affected by Asperger syndrome (i.e., high-functioning autism) or attention deficit hyperactivity disorder (ADHD). Here, we examine normative performance in a neurotypical population of young adults, therefore, all subjects filled out the Adult ADHD Self-Report Scale (ASRS, Kessler *et al*., 2005) and the autism-spectrum quotient (AQ, Baron-Cohen *et al*., 2001) questionnaires. The results from these questionnaires are summarized in Table 1. We subsequently calculated Pearson’s linear correlation coefficients between all tests and the ASRS and AQ scores. The correlation plots, including the correlation coefficients and p-values are shown in the supplementary ASRS_AQ_correlations.pdf. None of these correlations was significant in our neurotypical population.

## Discussion

The first goal of this study was to investigate the repeatability of the AIM and FInD paradigms in neurotypical healthy young adults for a variety of visual functions (acuity, contrast sensitivity, form- and motion coherence) performed on a tablet computer.

### Comparison with previous FInD and AIM studies

AIM Visual Acuity: Previous studies investigated the repeatability of AIM VA for a viewing distance of 4m instead of near testing as performed in the current study (Skerswetat *et al*., 2024a). That study showed that increasing the number of adaptive charts reduces the repeatability variance as expressed using limits of agreement which for 3 charts i.e., one initial and two adaptive steps ±0.24 logMAR was comparable to the ±0.14 logMAR (CoR: ±0.19 logMAR) of the current study. Also, the current study and the previous study found no systematic retest bias.

Form and Motion: A study in patients with and without cerebral visual impairment (Merabet *et al*., 2023) reported 27% for circular motion and 30% for form thresholds using the hidden cell FInD tasks in the control cohort, which is slightly different to 18% and 39%, respectively, as reported in the current study’s first test session (Table 2). This may be due to the different stimulus properties of the tablet used here and the 32” computer display used in Merabet *et al*. (2023). We are not aware of any published estimates of test-retest repeatability of coherence measured with other paradigms, such as 2AFC. In a study concerning vision in people with and without albinism (Skerswetat *et al*., 2024b), thresholds for the control group, using FInD circular Motion and Form Coherence with hidden cells mode, were 22% and 52%, respectively. Again, higher sensitivity in motion compared to form coherence was shown, but lower sensitivity for form coherence, which can be explained due to the higher age of the cohort since form coherence thresholds increase with age (McKendrick, Weymouth & Battista, 2013). FInD complex motion and form coherence tasks showed no significant mean biases, whereas bias was found here for translation, simple form and motion signals (Table 2). Future research may investigate whether the bias reflects learning and might disappear between a second and third session. In the meantime, it is suggested to run an initial training session prior to an actual task for these stimulus types. The study also revealed a wide inter-individual difference in the ability to perceive global motion and form (Figure 3).

AIM CSF 2D + showed no retest bias but had significantly larger variance when using three charts compared to five charts. Hence, it is suggested to use five charts as default to ensure minimum retest variance.

### Psychometric parameters

Some previous studies have investigated slope parameters (e.g., Wallis *et al*., 2013), however to our best knowledge no study specifically looked at the repeatability of slope parameters. Table 3 shows the slope and table 4 the minimum report error or *noise* (i.e., result of all noise factors) with the corresponding repeatability results.

**Table 3.**
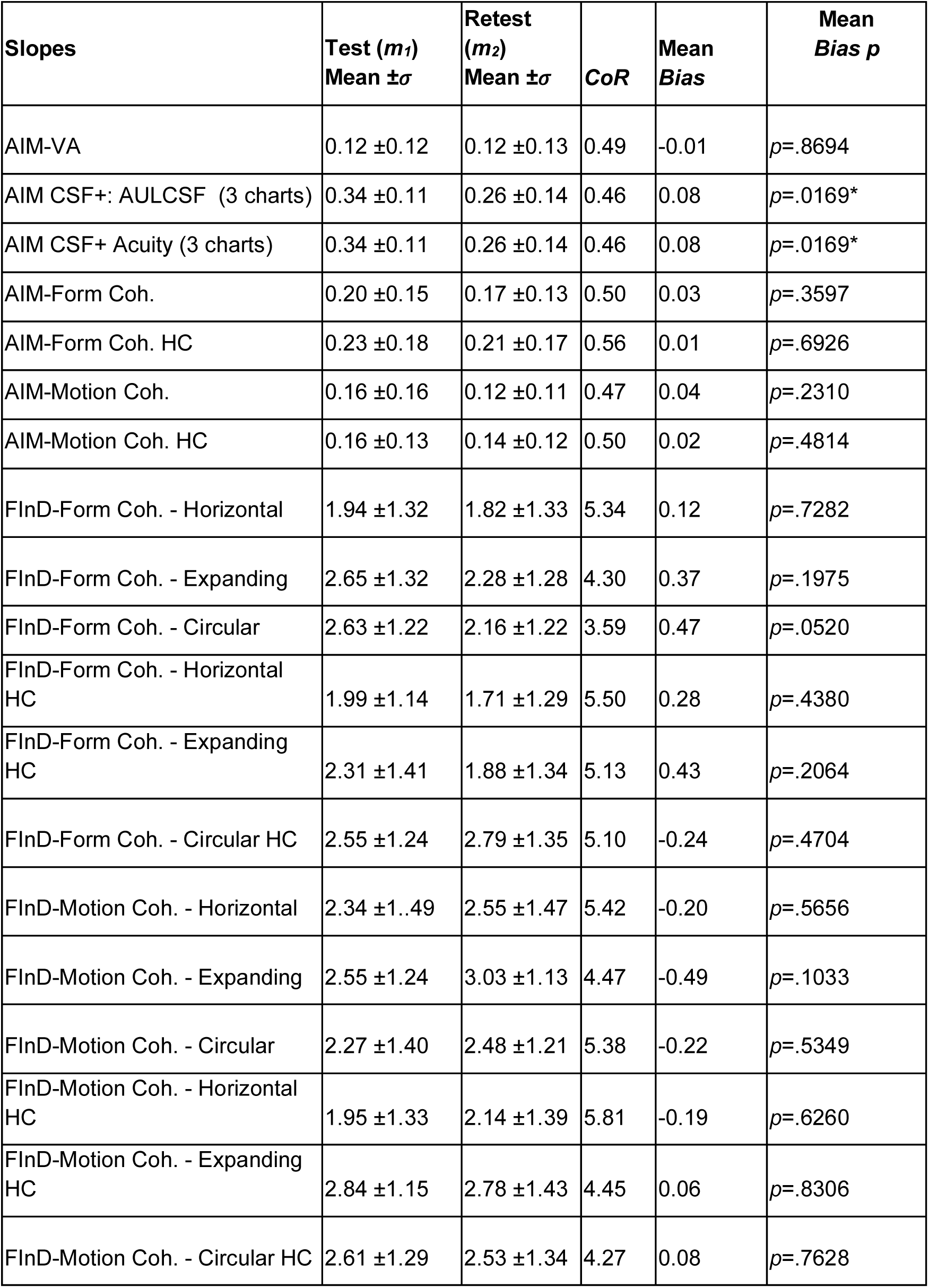
Test Retest results for slope values.

| **Slopes** | **Test ($m_1$)**<br>**Mean $\pm \sigma$** | **Retest ($m_2$)**<br>**Mean $\pm \sigma$** | **CoR** | **Mean**<br>**Bias** | **Mean**<br>**Bias $p$** |
| --- | --- | --- | --- | --- | --- |
| AIM-VA | 0.12 ± 0.12 | 0.12 ± 0.13 | 0.49 | -0.01 | $p=.8694$ |
| AIM CSF+: AULCSF (3 charts) | 0.34 ± 0.11 | 0.26 ± 0.14 | 0.46 | 0.08 | $p=.0169^*$ |
| AIM CSF+ Acuity (3 charts) | 0.34 ± 0.11 | 0.26 ± 0.14 | 0.46 | 0.08 | $p=.0169^*$ |
| AIM-Form Coh. | 0.20 ± 0.15 | 0.17 ± 0.13 | 0.50 | 0.03 | $p=.3597$ |
| AIM-Form Coh. HC | 0.23 ± 0.18 | 0.21 ± 0.17 | 0.56 | 0.01 | $p=.6926$ |
| AIM-Motion Coh. | 0.16 ± 0.16 | 0.12 ± 0.11 | 0.47 | 0.04 | $p=.2310$ |
| AIM-Motion Coh. HC | 0.16 ± 0.13 | 0.14 ± 0.12 | 0.50 | 0.02 | $p=.4814$ |
| FInD-Form Coh. - Horizontal | 1.94 ± 1.32 | 1.82 ± 1.33 | 5.34 | 0.12 | $p=.7282$ |
| FInD-Form Coh. - Expanding | 2.65 ± 1.32 | 2.28 ± 1.28 | 4.30 | 0.37 | $p=.1975$ |
| FInD-Form Coh. - Circular | 2.63 ± 1.22 | 2.16 ± 1.22 | 3.59 | 0.47 | $p=.0520$ |
| FInD-Form Coh. - Horizontal HC | 1.99 ± 1.14 | 1.71 ± 1.29 | 5.50 | 0.28 | $p=.4380$ |
| FInD-Form Coh. - Expanding HC | 2.31 ± 1.41 | 1.88 ± 1.34 | 5.13 | 0.43 | $p=.2064$ |
| FInD-Form Coh. - Circular HC | 2.55 ± 1.24 | 2.79 ± 1.35 | 5.10 | -0.24 | $p=.4704$ |
| FInD-Motion Coh. - Horizontal | 2.34 ± 1.49 | 2.55 ± 1.47 | 5.42 | -0.20 | $p=.5656$ |
| FInD-Motion Coh. - Expanding | 2.55 ± 1.24 | 3.03 ± 1.13 | 4.47 | -0.49 | $p=.1033$ |
| FInD-Motion Coh. - Circular | 2.27 ± 1.40 | 2.48 ± 1.21 | 5.38 | -0.22 | $p=.5349$ |
| FInD-Motion Coh. - Horizontal HC | 1.95 ± 1.33 | 2.14 ± 1.39 | 5.81 | -0.19 | $p=.6260$ |
| FInD-Motion Coh. - Expanding HC | 2.84 ± 1.15 | 2.78 ± 1.43 | 4.45 | 0.06 | $p=.8306$ |
| FInD-Motion Coh. - Circular HC | 2.61 ± 1.29 | 2.53 ± 1.34 | 4.27 | 0.08 | $p=.7628$ |

**Table 4.** Test-retest results for minimum report errors values.

| Errors [°] | Test ($m_1$)<br>Mean $\pm\sigma$ | Retest ($m_2$)<br>Mean $\pm\sigma$ | CoR | Mean<br>Bias | Mean<br>Bias $p$ |
| --- | --- | --- | --- | --- | --- |
| AIM-VA | 12.05 ±5.77 | 11.43 ±5.65 | 22.27 | 0.62 | $p=.6695$ |
| AIM CSF+: AULCSF | 8.32 ±3.14 | 8.41 ±3.01 | 9.96 | -0.09 | $p=.8900$ |
| AIM CSF+ Acuity | 8.32 ±3.14 | 8.41 ±3.01 | 9.96 | -0.09 | $p=.8900$ |
| AIM-Form Coh. | 10.10 ±6.74 | 12.36 ±7.58 | 23.20 | -2.26 | $p=.1430$ |
| AIM-Form Coh. HC | 9.41 ±7.87 | 11.14 ±8.89 | 27.64 | -1.73 | $p=.3410$ |
| AIM-Motion Coh. | 15.94 ±15.71 | 15.40 ±15.80 | 47.84 | 0.54 | $p=.8631$ |
| AIM-Motion Coh. HC | 16.29 ±13.39 | 13.25 ±15.78 | 53.69 | 3.04 | $p=.3894$ |
| FInD-Form Coh. - Horizontal | 0.09 ±0.11 | 0.07 ±0.09 | 0.33 | 0.01 | $p=.5649$ |
| FInD-Form Coh. - Expanding | 0.08 ±0.09 | 0.09 ±0.13 | 0.33 | 0.00 | $p=.8917$ |
| FInD-Form Coh. - Circular | 0.04 ±0.07 | 0.07 ±0.08 | 0.15 | -0.03 | $p=.0097^*$ |
| FInD-Form Coh. - Horizontal<br>HC | 0.12 ±0.14 | 0.08 ±0.11 | 0.38 | 0.04 | $p=.0896$ |
| FInD-Form Coh. - Expanding<br>HC | 0.14 ±0.18 | 0.07 ±0.08 | 0.49 | 0.07 | $p=.0336^*$ |
| FInD-Form Coh. - Circular HC | 0.08 ±0.10 | 0.07 ±0.10 | 0.29 | 0.01 | $p=.4588$ |
| FInD-Motion Coh. - Horizontal | 0.13 ±0.17 | 0.12 ±0.17 | 0.48 | 0.01 | $p=.8697$ |
| FInD-Motion Coh. - Expanding | 0.05 ±0.07 | 0.03 ±0.06 | 0.23 | 0.02 | $p=.2093$ |
| FInD-Motion Coh. - Circular | 0.04 ±0.07 | 0.07 ±0.14 | 0.24 | -0.03 | $p=.0982$ |
| FInD-Motion Coh. - Horizontal<br>HC | 0.13 ±0.20 | 0.13 ±0.16 | 0.50 | 0.00 | $p=.8969$ |
| FInD-Motion Coh. - Expanding<br>HC | 0.08 ±0.09 | 0.08 ±0.13 | 0.29 | 0.00 | $p=.8124$ |
| FInD-Motion Coh. - Circular<br>HC | 0.03 ±0.04 | 0.05 ±0.13 | 0.33 | -0.02 | $p=.3096$ |

With the exception of CSF results using three charts, all slopes were bias free. The noise parameters were mostly bias-free except for FInD Form circular coherence and hidden cell expanding coherence. These results show that the methods employed in the AIM and FInd paradigms produce consistent and repeatable measurements.

### Testing durations

The average time for visual acuity (Table 5) was less than two minutes in total and an approximate chart duration of 30 seconds, which is comparable to previously reported times when testing for distance visual acuity (Skerswetat *et al*., 2024a). The mean chart duration was less than one minute for the CSF task, hence the total duration for 5 charts is approximately 4 minutes, which is faster than conventional tests that may take 25 minutes (Kelly & Savoie, 1973) and comparable to other adaptive rapid procedures (Liu *et al*., 2021).

**Table 5.** Summary of overall testing time and per-chart time.

| Visual function test | Total Duration [s] |  | Chart duration [s] |  |
| --- | --- | --- | --- | --- |
| | Test ($m_1$)<br>Mean $\pm \sigma$ | Retest ($m_2$)<br>Mean $\pm \sigma$ | Test ($m_1$)<br>Mean $\pm \sigma$ | Retest ($m_2$)<br>Mean $\pm \sigma$ |
| AIM-Visual Acuity | 109 ±40 | 92 ±34 | 36 ±13 | 31 ±11 |
| AIM CSF+ (3 charts) | 145 ±64 | 111 ±46 | 48 ±21 | 37 ±15 |
| AIM- Form Coherence | 196 ±111 | 177 ±21 | 65 ±40 | 59 ±82 |
| AIM- Form Coherence HC | 244 ±191 | 164 ±107 | 81 ±64 | 55 ±36 |
| AIM- Motion Coherence | 222 ±90 | 176 ±34 | 74 ±30 | 59 ±34 |
| AIM- Motion Coherence HC | 283 ±126 | 216 ±83 | 94 ±42 | 72 ±28 |
| FInD- Form Coherence | 263 ±288 | 182 ±124 | 29 ±32 | 20 ±14 |
| FInD- Form Coherence HC | 365 ±332 | 225 ±140 | 40 ±37 | 25 ±16 |
| FInD- Form Coherence | 306 ±165 | 248 ±99 | 34 ±18 | 28 ±11 |
| FInD- Form Coherence HC) | 360 ±139 | 301 ±149 | 40 ±15 | 33 ±17 |

The test durations for the Form and Motion tasks varied more widely specifically between 3 and 6 minutes per tests, one because the FInD task tested three subtypes of form and motion percepts, namely horizontal, circular and expanding percepts, compared to translational, simple form or motion percepts. The hidden cell paradigms caused longer testing durations than the open cell paradigms. This may be due to either the use of a mouse instead of finger touches, different gaze strategies used to detect signals, or a combination of those reasons. Since we did not collect gaze position data, future studies may investigate whether different gaze strategies were used for open and hidden cell modes.

### Functional relationship between psychometric outcomes

The second goal of the study was to investigate relationships among thresholds. We used a combination of linear models and hierarchical cluster analysis, unlike in a previous study where these analyses were performed separately (Skerswetat *et al*., 2024b). The current study shows overall within-function outcomes clustering, specifically for contrast - contrast, form - form, and motion - motion outcomes (Figure 2). Similarly, contrast outcomes correlated with each other, as well as motion outcomes. Skerswetat *et al*. (2024b) and Bosten *et al*. (2017) also found significant correlations among contrast outcomes, but not among motion values. We speculate that this difference may be due to the more heterogeneous age range and smaller sample within the control cohort of that study compared to the current study. The same study (Skerswetat *et al*., 2024b) also showed clustering within contrast and within form and motion outcomes for the control cohort, which is in agreement with the current study. Overall, these results suggest distinct processing pathways of contrast and mid-level form and motion perception.

Although the primary aim of this study was to assess the repeatability of the tests, the results—particularly those obtained from the AIM and FInD Form and Motion tests—also provide insight into the visual processing of global texture and motion. A detailed analysis, along with further discussion and comparisons with previous studies, is presented in the Appendix and Figure A2.

### Future directions

The current study investigated the repeatability and functional relationship of the tablet-based psychophysical tests in young healthy participants. Based on the high repeatability of these results and the normative data, future research can be adequately powered to test the effect of age and the effect in clinically relevant cohorts on the test performance. The number of cells within each chart and the number of charts determine the testing duration, and the present data can guide future research to investigate their relationship to optimise the time-repeatability tradeoff.

## Conclusion

AIM and FInD near vision tests are generalizable across multiple visual functions and are precise and reliable. Most functions tested were bias-free, except translational form and motion coherence, which showed significant improvement on a second test. In future studies, such bias may be improved with additional instructions or training charts. CSF, form, and motion outcomes clustered together, and CSF and motion outcomes correlated with one another, indicating separate processing mechanisms for achromatic spatial and global coherence perception.

Together, the results demonstrate the reliability of the tests, which may be useful for basic and clinical research.

## Acknowledgements

Conflict of interest: JS and PJB are co-founders and shareholders of PerZeption Inc. JS and PJB are the inventors of AIM and FInD technology (P). Northeastern University is the owner of the intellectual property and granted exclusive licensing rights to PerZeption Inc.

GS and JG were supported by the College of Optometrists (UK).

PJB and JS were supported by NIH R01EY032162.

## Appendix

**Figure A1.**
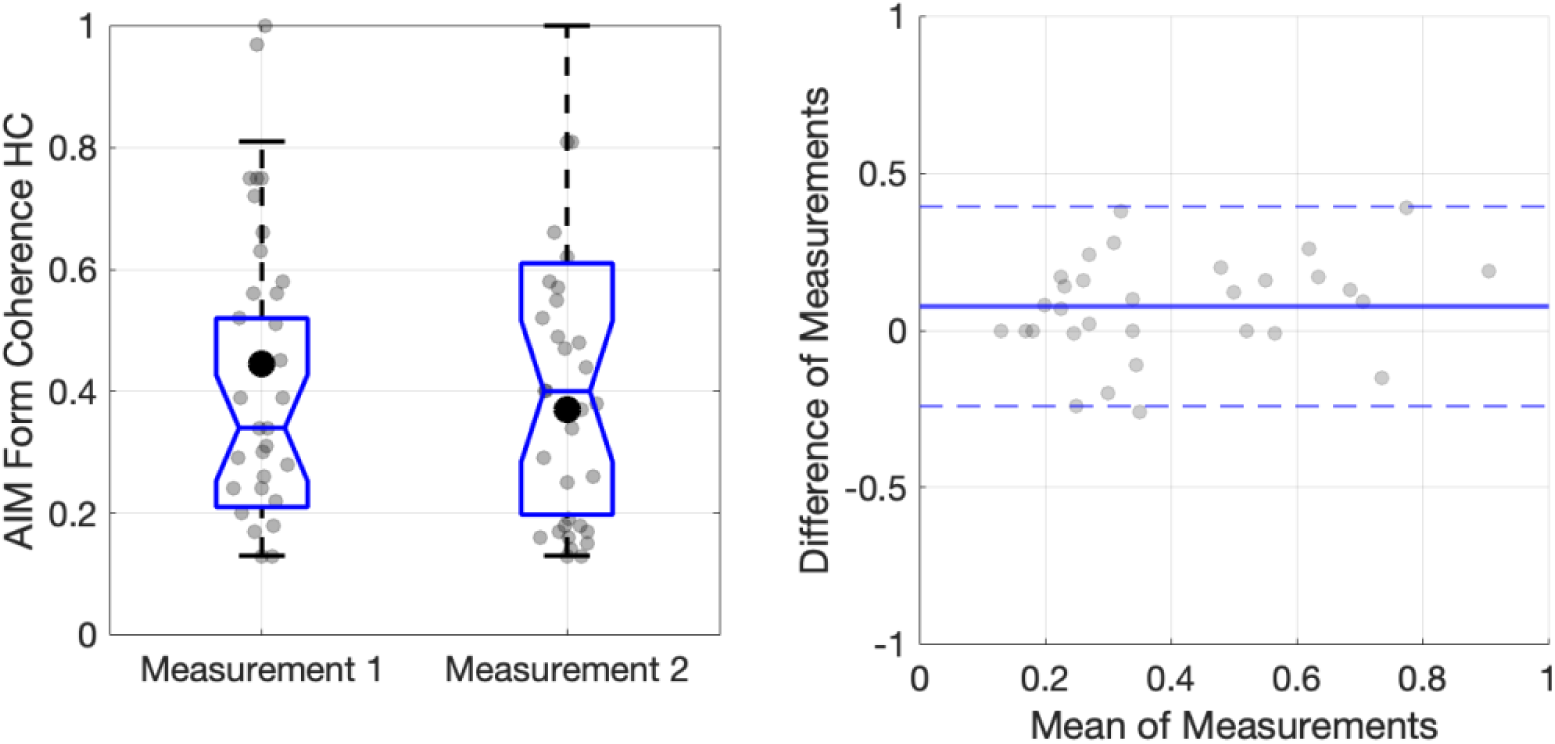
Left: Example data showing Box and Whisker plots with individual data (grey data points) and the mean (black solid data point) for the AIM Form Coherence test (coherence thresholds are expressed in %, 0-1). Right: Bland-Altman plots for AIM Form Coherence. Figures for all tests can be found in the supplementary material (Bland_Altman_plots.pdf).

### Comparison between horizontal, expanding and circular form and motion

The results from the AIM and FInD Form and Motion tests have implications regarding the visual processing of global texture and motion. There is ample psychophysical and physiological evidence for the hypothesis that the representation of objects and shapes is achieved by a hierarchical feedforward process along the ventral pathway (e.g., Cadieu *et al*., 2007; Serre *et al*., 2005; Serre, Oliva, & Poggio, 2007; Van Essen, Anderson, & Felleman, 1992). A key question in vision science is how local information detected in the early visual areas (primary visual cortex) is integrated to represent more complex, behaviorally relevant stimuli (e.g., faces, objects) at subsequent stages. Psychophysically, this has been tested in different ways. One approach, also employed in the Form and Motion tests, is to determine the minimum number of signal elements required to detect the presence of a global pattern embedded in an array of noise elements (coherence thresholds). This measurement of coherence thresholds has been used in many studies of motion (Braddick *et al*., 2000; Newsome, 1988), texture (Dakin, 1997; Schmidtmann *et al*., 2015; Wilson & Wilkinson, 1998; Wilson, Wilkinson, & Asaad, 1997), and contour perception (Achtman, Hess, & Wang, 2003; Braddick *et al*., 2000; Schmidtmann *et al*., 2013; 2015). Coherence thresholds in Glass patterns (Glass, 1969; Glass & Perez, 1973) have been employed to investigate signal integration in texture perception (Dakin, 1997; Dakin & Bex, 2002; Dickinson, Broderick, & Badcock, 2009; Wilson, Wilkinson & Asaad, 1997; Wilson & Wilkinson, 1998). Models of Glass pattern detection propose a two-stage process, in which the first stage extracts local orientation information while the second stage integrates the orientation information in order to extract the global structure (Wilson, Wilkinson & Assad, 1997; Wilson & Wilkinson, 1998, see Schmidtmann *et al*. (2015) for a revised model). This model is supported by physiological data showing that neither the classical receptive fields nor surround mechanisms in early visual areas (V1, V2) are sufficient to process the Glass patterns (Smith, Bair, & Movshon, 2002; Smith *et al*., 2007). Some previous psychophysical studies found no difference in sensitivities for detecting circular and radial structure (e.g., see Kelly *et al*., 2001; Kurki & Saarinen, 2004; Pei *et al*., 2005; Seu & Ferrera, 2001; Wilson, Wilkinson & Asaad, 1997; Wilson & Wilkinson, 1998), while others reported a significantly increased sensitivity of human observers for circular, followed by radial, spiral, and parallel structure (Kelly *et al*., 2001; Kurki & Saarinen, 2004; Seu & Ferrera, 2001; Wilson, Wilkinson & Asaad, 1997; Wilson & Wilkinson, 1998), suggesting that the global structure in Glass patterns is processed by specialized detectors for circular and radial textures. However, there is also contradictory evidence. For instance, Dakin and Bex (2002) found that the advantage for circular Glass patterns was only evident if the stimulus was presented within a circular window as opposed to a square window. Similarly, Schmidtmann *et al*. (2015) were not able to replicate the above mentioned hierarchy. Their results showed that detection sensitivity was independent of texture type; hence, no evidence for specialized detectors for circular structure.

The results presented here also do not show the clear sensitivity hierarchy (i.e., circular > expanding > horizontal) described above. Figure A2 shows Box and Whisker plots comparing coherence thresholds achieved for horizontal, expanding and circular structure for the FInD Form, FInD Form HC, FInD Motion, and FInD Motion HC tests. Following Shapiro-Wilk tests for normality, which showed that some of the conditions were not normally distributed, the data were analyzed using a non-parametric Friedman ANOVA (Durbin-Conover) across all conditions. This analysis revealed a statistically significant global effect (𝝌^2^(119) = 11, p<.001). Subsequent pairwise comparisons (Bonferroni-corrected for multiple comparisons) showed that for the FInD Form paradigms, sensitivity was largely independent of texture type, except for a statistically significant increase in thresholds (i.e. decrease in sensitivity) between FInD Form HC horizontal and FInD Form HC circular. In general, performance for the FInD Motion paradigms was slightly (sometimes significantly better; see Table A1) compared to the static FInD Form stimuli. Results show that sensitivity for FInD Motion horizontal and FInD Motion expanding stimuli was significantly lower compared to FInD Motion circular. Finally, sensitivity for FInD Motion HC expanding was significantly lower compared to FInD Motion HC circular and FInD Motion HC horizontal. No superior performance for circular motion is evident.

**Figure A2.**
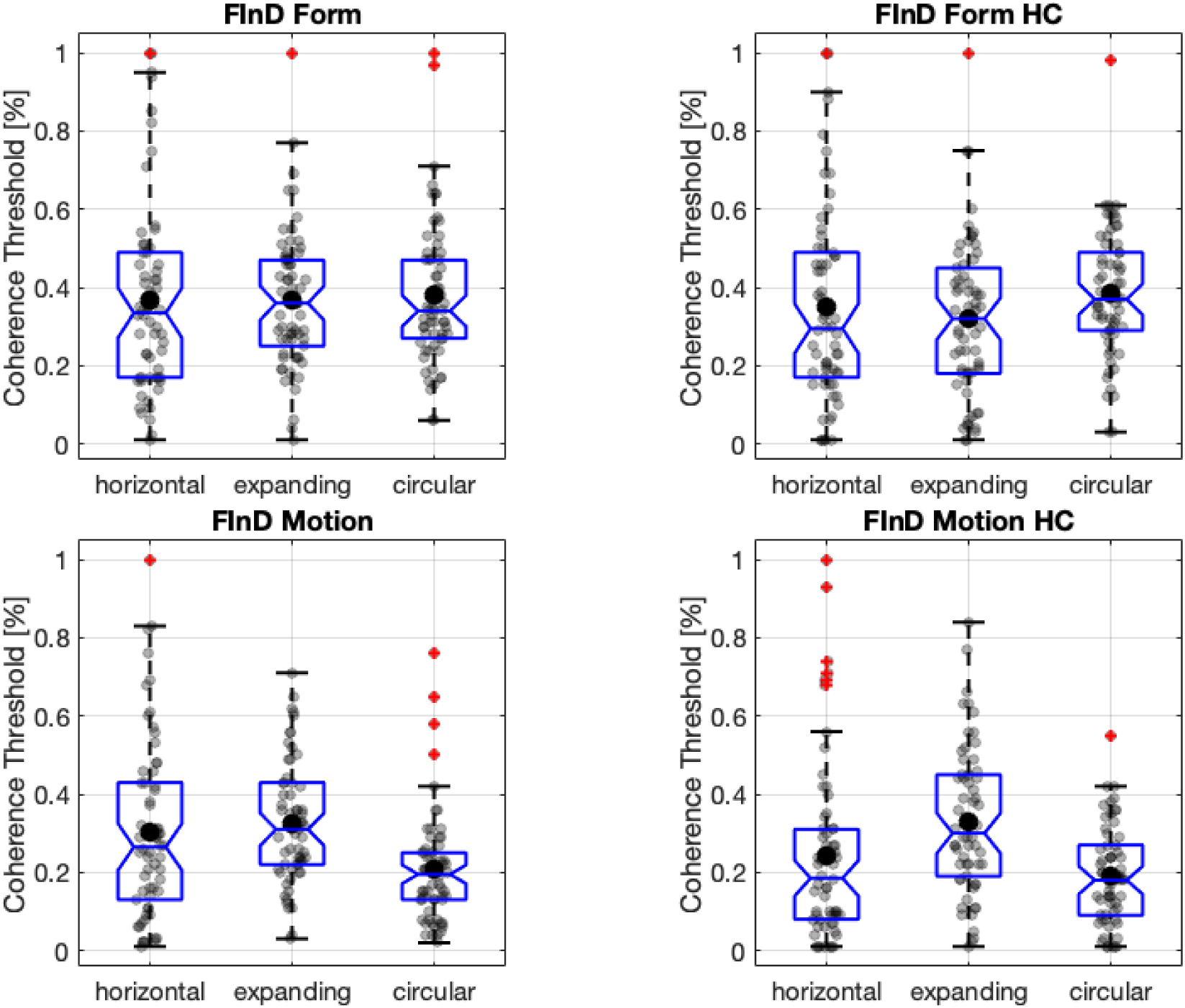
Box and Whisker plots with individual data (grey data points) and the mean (black solid data point) for the FInD Form (top left), FInD Form HC (top right), FInD Motion (bottom left) and FInD Motion HC (bottom right) tests (coherence thresholds are expressed in %, 0-1). The outliers (red diamonds) with coherence thresholds of ∼1 (∼100%) were from two subjects who could not do the task, independent of global structure.

**Table A1.**
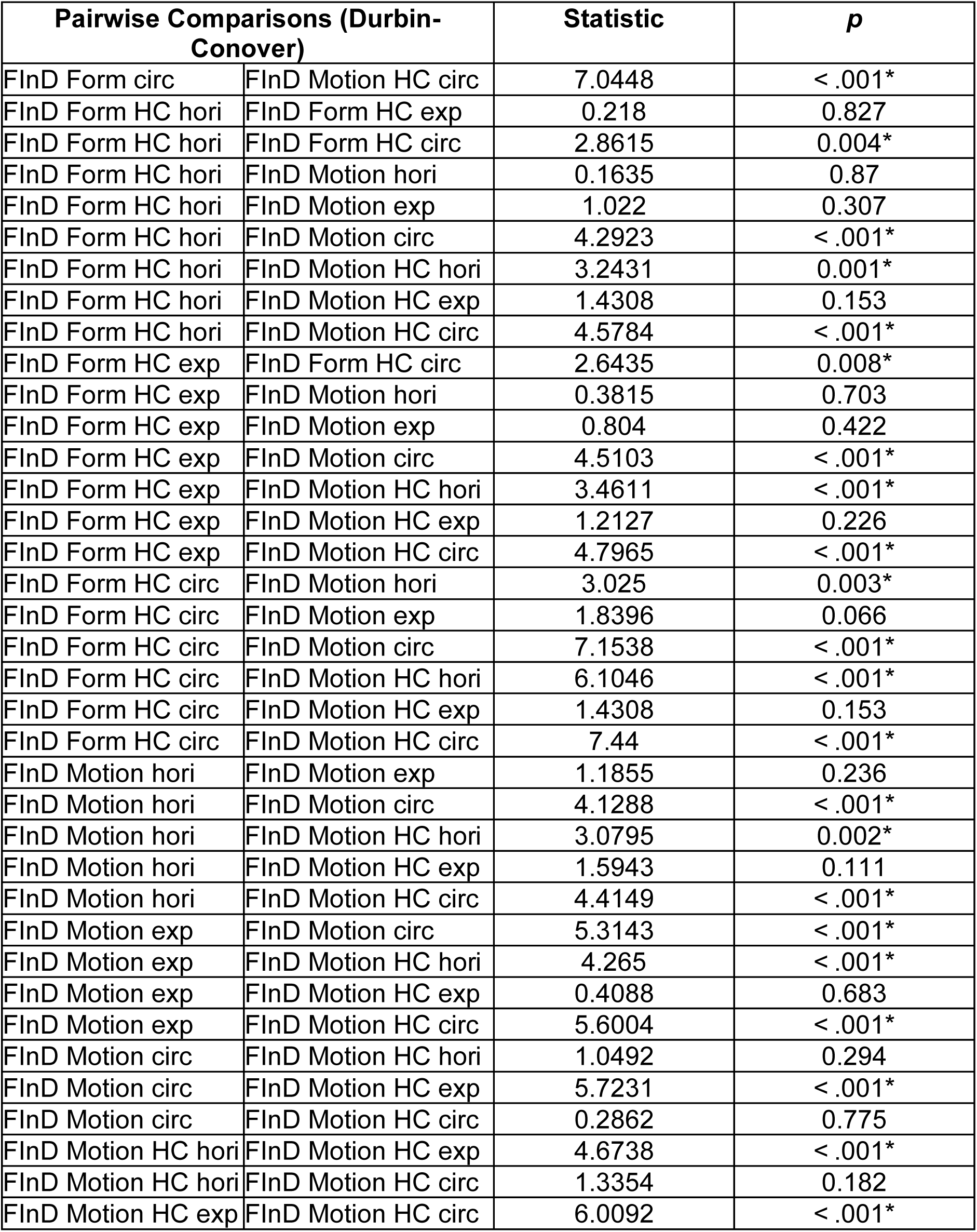
non-parametric Friedman ANOVA pairwise comparison (Durbin-Conover). Statistically significant conditions are indicated by *.

| Pairwise Comparisons (Durbin-Conover) |  | Statistic | *p* |
| --- | --- | --- | --- |
| FInD Form circ | FInD Motion HC circ | 7.0448 | < .001* |
| FInD Form HC hori | FInD Form HC exp | 0.218 | 0.827 |
| FInD Form HC hori | FInD Form HC circ | 2.8615 | 0.004* |
| FInD Form HC hori | FInD Motion hori | 0.1635 | 0.87 |
| FInD Form HC hori | FInD Motion exp | 1.022 | 0.307 |
| FInD Form HC hori | FInD Motion circ | 4.2923 | < .001* |
| FInD Form HC hori | FInD Motion HC hori | 3.2431 | 0.001* |
| FInD Form HC hori | FInD Motion HC exp | 1.4308 | 0.153 |
| FInD Form HC hori | FInD Motion HC circ | 4.5784 | < .001* |
| FInD Form HC exp | FInD Form HC circ | 2.6435 | 0.008* |
| FInD Form HC exp | FInD Motion hori | 0.3815 | 0.703 |
| FInD Form HC exp | FInD Motion exp | 0.804 | 0.422 |
| FInD Form HC exp | FInD Motion circ | 4.5103 | < .001* |
| FInD Form HC exp | FInD Motion HC hori | 3.4611 | < .001* |
| FInD Form HC exp | FInD Motion HC exp | 1.2127 | 0.226 |
| FInD Form HC exp | FInD Motion HC circ | 4.7965 | < .001* |
| FInD Form HC circ | FInD Motion hori | 3.025 | 0.003* |
| FInD Form HC circ | FInD Motion exp | 1.8396 | 0.066 |
| FInD Form HC circ | FInD Motion circ | 7.1538 | < .001* |
| FInD Form HC circ | FInD Motion HC hori | 6.1046 | < .001* |
| FInD Form HC circ | FInD Motion HC exp | 1.4308 | 0.153 |
| FInD Form HC circ | FInD Motion HC circ | 7.44 | < .001* |
| FInD Motion hori | FInD Motion exp | 1.1855 | 0.236 |
| FInD Motion hori | FInD Motion circ | 4.1288 | < .001* |
| FInD Motion hori | FInD Motion HC hori | 3.0795 | 0.002* |
| FInD Motion hori | FInD Motion HC exp | 1.5943 | 0.111 |
| FInD Motion hori | FInD Motion HC circ | 4.4149 | < .001* |
| FInD Motion exp | FInD Motion circ | 5.3143 | < .001* |
| FInD Motion exp | FInD Motion HC hori | 4.265 | < .001* |
| FInD Motion exp | FInD Motion HC exp | 0.4088 | 0.683 |
| FInD Motion exp | FInD Motion HC circ | 5.6004 | < .001* |
| FInD Motion circ | FInD Motion HC hori | 1.0492 | 0.294 |
| FInD Motion circ | FInD Motion HC exp | 5.7231 | < .001* |
| FInD Motion circ | FInD Motion HC circ | 0.2862 | 0.775 |
| FInD Motion HC hori | FInD Motion HC exp | 4.6738 | < .001* |
| FInD Motion HC hori | FInD Motion HC circ | 1.3354 | 0.182 |
| FInD Motion HC exp | FInD Motion HC circ | 6.0092 | < .001* |

